# Primary cilia sensitize insulin receptor-mediated negative feedback in pancreatic β cells

**DOI:** 10.1101/2020.12.02.408914

**Authors:** Yuezhe Li, Prem. K. Shrestha, Yi I. Wu

## Abstract

Insulin receptors (IR) can localize to the primary cilia of pancreatic β cells. Because primary cilia are known to sensitize or bias the signaling of cell surface receptors, we investigate how ciliary insulin receptors influence glucose-stimulated insulin secretion (GSIS) in β cells by gauging how cytosolic calcium concentration changes in a mouse insulinoma cell line (MIN6). Purified recombinant insulin suppresses calcium elevation in response to glucose in these cells. Interestingly, ciliated cells show attenuated cytosolic calcium elevation compared to cilium-free cells after glucose stimulation even in the absence of exogenous insulin. We observe that ciliary IR is highly phosphorylated, and the phospho-IR density decreases when cells are either treated with an insulin receptor (IR) inhibitor, BMS536924, or ciliary function is disrupted through either IFT88 or BBS1 knockdown. Consistently, the attenuation of calcium elevation in ciliated cells is abrogated when cells are either treated with IR inhibitor or when primary cilia are impaired. We further demonstrate that ciliary IR signaling hyperpolarizes the plasma membrane but has no apparent impact on glucose-induced ATP production. Thus, our results argue that primary cilia sensitize insulin receptor signaling and mediate negative feedback in GSIS in pancreatic β cells.

## Introduction

Pancreatic beta cells (β cells) are responsible for secreting insulin and maintain energy homeostasis in the body. Upon elevation of blood glucose, β cells accelerate glucose metabolism and increase the production of ATP. The primary outcome of such changes is the increase of ATP/ADP ratio in the cytosol, which causes the closure of ATP-sensitive potassium channels and membrane depolarization. The subsequent opening of the voltage-dependent calcium channels leads to the elevation of cytosolic calcium, which triggers insulin exocytosis (MacDonald et al., 2005). How β cells sense and launch suitable responses to glucose change remains an active area of diabetes research.

Insulin feedback are thought to regulate β cell function. Insulin signals through either the insulin receptor (IR) or the insulin-like growth factor 1 receptor (IGF1R) (White, 2003). In addition, there are two isoforms of IR, insulin receptor A (IR-A) and insulin receptor B (IR-B), which differ by 12 amino acids (Seino et al., 1990) but signal differently (Leibiger et al., 2001). All of these receptors are expressed in pancreatic β cells, and their signaling are important for maintaining β cell mass, growth, insulin synthesis, and secretion in the long run (reviewed in Leibiger et al., 2008). However, the acute effects of insulin receptor signaling on secretion are more elusive. Some studies report that insulin or insulin mimetics suppresses insulin secretion in humans, mice or in isolated islets (Elahi et al., 1982, Argoud et al., 1987, Persaud et al., 2002, Zhang et al., 1999). Phosphoinositide 3-kinase (PI3K)-dependent activation of the ATP-sensitive potassium channels are thought to mediate this effect (Khan et al., 2001). Openings ATP-sensitive potassium channels lead to membrane hyperpolarization, thus inhibiting the electrical activity and calcium elevation of β cells. However, other studies report insulin or insulin mimetics either have dual effects on insulin secretion (Jimenez-Feltstrom et al., 2004) or promote exocytosis (Roper et al., 2002, Bouche et al., 2010, Aoyagi et al., 2010, Aspinwall et al., 1999, Aspinwall et al., 2000, Hisanaga et al., 2009). The stimulatory effects of insulin on insulin secretion is also dependent on PI3K activity. IR signaling elevates cytosolic calcium through either inhibiting calcium sequestering into the endoplasmic reticulum (ER) or through the translocation of transient receptor potential cation channel subfamily V member 2 (TRPV2), a calcium-permeable channel (Ramsey et al., 2006), from cytosol to the plasma membrane.

Recently, primary cilium has attracted much interest in diabetes research due to the strong association between type 2 diabetes and ciliopathies. Ciliopathies refer to a collection of several rare disorders linked to mutations in primary cilium proteins, such as Alstrom syndrome and Bardet-Biedl syndrome. Patients of Alstrom or Bardet-Biedl syndrome have frequent (~50 or greater) onsets of type 2 diabetes (diIorio et al., 2014). Studies in mice also linked ciliary dysfunction to abnormal insulin secretion (Gerdes et al., 2014, Zhang et al., 2005). Most if not all pancreatic β cells possess a primary cilium *in vivo* (Gan et al., 2017). Interestingly, IR-A was found enriched in the primary cilium immediately after insulin stimulation (Gerdes et al., 2014). Previous studies have shown that primary cilia can potentiate or suppress the signaling of their resident receptors, or modulate receptor signaling by influencing the preference of its downstream effectors (reviewed in Schou et al., 2015, Christensen et al., 2011). This has led us to hypothesize that ciliary insulin receptor may acutely affect the function of pancreatic β-cells.

In this study, we aim to find out the impact of IR signaling on GSIS and the role of primary cilium in modulating IR signaling. We demonstrate that insulin has a negative impact on glucose-induced calcium elevation in a β cell line in culture. We further demonstrate that primary cilium potentiates this effect by phosphorylating insulin receptor at residue concentration of insulin.

## Material and Methods

### Cell culture and transfection

MIN6 cells, a mouse insulinoma β cell line, were obtained from Dr. Jun-ichi Muyazaki. The cells were cultured in DMEM (Lonza, Basel, Switzerland), supplemented with 15% (v/v) fetal bovine serum (Gibco, Billings, MO), Penicillin/Streptomycin (Lonza), and 2.5 ul of BME as previously described (REF PMID 2163307) and were maintained under standard cell culture conditions (37 °C and 5% CO2). The cell lines were regularly checked for mycoplasma contamination. GeneCeillin DNA transfection reagent (Bulldog Bio, Portsmouth, NH) was used for transient transfections according to the manufacturer’s instructions. Lipofectamine 3000 (Thermo Fisher, Waltham, MA) was used to transfect siRNA in cells according to manufacturers’ protocol.

### DNA plasmids

LentiCas9-Blast (Addgene plasmid # 52962) was a gift from Dr. Feng Zhang (Massachusetts Institute of Technology, Cambridge). Arl13b was a gift from Dr. Takanari Inoue (John Hopkins University, Baltimore, MD). R-GECO1.2 (Addgee plasmid #45494) was a gift from Dr. Robert Campbell (University of Alberta, Edmonton, Alberta Canada). Insulin-GFP was a gift from Dr. Peter Arvan (University of Michigan, Ann Arbor). PercevalHR (Addgene plasmid #49083) is a gift from Dr. Gary Yellen (Harvard Medical School, Boston, U.S.A). Additional point mutations in the Arl13b, K216E and R219E, which reduce cytotoxicity, were generated using overlapping polymerase chain reaction (PCR). For transient transfection, the Arl13b mutant was subcloned into pmScarlet-I-N1 (modified from Addgene plasmid # 85044) or pmCherry-N1 (Clontech) vectors. For lentiviral constructs, R-GECO and Arl13b-mScarlet was generated by replacing the Cas9 gene in the LentiCas9-Blast backbone using restriction digestion or Gibson Assembly. The open reading frames of all DNA plasmids were verified by DNA sequencing.

### RNAi approaches

siGENOME non-targeting (D-001206-13-05, D-001210-04-05), IFT88 (M-050417-00-0005), BBS1(M-056626-01-0005) siRNA, IR (M-043748-01-0005) siRNA pools (Horizon, Dharmacon, Colorado, USA) were used according to the manufacturer’s instructions.

### Western blot

MIN6 cells prior and after the addition of either bovine insulin, recombinant human insulin, or recombinant human IGF1 were washed three times with PBS and lysed on ice in a buffer containing 0.1% Triton X-100, 50 mM monobasic sodium phosphate (pH 7.4), 150 mM NaCl, 5 mM EDTA and Halt Protease inhibitor cocktail (Thermo Scientific, Rockford, IL, USA). The cell lysates were fractionated using 4–12% NuPAGE gels (Invitrogen), immobilized on to PVDF membranes (Millipore, Germany), followed by immunoblotting using anti-phospho-IR/IGF1R polyclonal antibody (19H7, Cell Signaling, Danvers, MA, USA), anti-insulin receptor β polyclonal antibody (4B8, Cell Signaling, Danvers, MA, USA), anti-IFT88 polyclonal antibody (Abcam, Cambridge, UK), anti-BBS1 polyclonal antibody (Proteintech, Rosemont, IL, USA), or anti-GAPDH monoclonal antibody (Millipore, Germany) and horseradish peroxidase-coupled secondary antibody (Millipore). Enhanced chemiluminescence detection (Pierce, Waltham, MA, USA) and ChemiDoc MP Imaging System (Bio-Rad, Hercules, CA, U.S.A) were used to detect the signals. For the purpose of clarity, the blot images were cropped to show relevant bands.

### Immunofluorescence

MIN6 cells were seeded on fibronectin-coated cover glasses 3 days prior to experiments. Plasmids/ siRNA were transfected 2 days prior to experiments. The cells were fixed in 4% formaldehyde for 20 minutes before being permeabilized in 0.3% Triton X for 20 minutes. Cells were then blocked in 1% BSA for 20 minutes before being incubated in a primary antibody against acetylated α tubulin (1:4000 dilution, clone 6-11B-1, Sigma-Aldrich, St. Louis, MO, USA), GFP (1:1000, clone A11120, Thermo Fisher, Waltham, MA), Arl13b (1:500, Proteintech, Rosemont, IL, USA), phospho-IR/ IGF1R (Tyr1162, Tyr1163) (1:100, Thermo Fisher, Waltham, MA) either at 4°C overnight or at room temperature for 45 minutes or 3 hours. Cells were incubated in secondary antibodies (1:1000 dilution) for 45 minutes at room temperature. Cell nuclei were labeled with DAPI (0.2mg/ml, Thermo Fisher, Waltham, MA) for 3 minutes.

### Imaging setup

All time-lapse imaging experiments were performed on a customized Nikon Ti-E inverted microscope. To improve signal noise ratio, total internal reflection fluorescence microscopy (TIRFM) was performed using a 60x oil TIRF objective (NA 1.49). The penetration depth of the evanescent waves from different laser lines were calibrated as described previously (Ref PMID 19816922). The microscope was modified with a “stage-up” design which enables the insertion of two independent, motorized dichroic mirrors/filter cubes in the microscope infinity space. A dichroic mirror in the bottom cube was used to reflect excitation laser lines at 488, 561, 640 nm (Coherent OBIS) for imaging of Cal520 (AAT Bioquest, Sunnyvake, CA), R-GECO, Di-4-ANEQ(F)PTEA (Potentioetric Probe, Farmington, CT), respectively. The fluorescent emission was captured with an EMCCD camera (iXon Ultra, Andor). Live cell imaging was performed at 37°C in a heated chamber (Bioptechs) with a humidified 5% CO2 supply. Customized vitamin and phenol red-free DMEM medium (US Biological) was used in live cell imaging to reduce fluorescence background and photobleaching. For imaging of immunofluorescence labeled cells, a Nikon TE2000 microscope equipped with a CSU-10 spinning disk confocal unit (Yokogawa) was used. The laser excitation light sources were from Coherent OBIS (405nm) and Melles Griot (488, 568 and 647nm). Metamorph software (Molecular Devices) was used to control both imaging setups.

### Measurement and analysis of cytosolic calcium concentration

The calcium fluctuations in MIN6 cells were tracked using either R-GECO (lentivirally tranduced stable cell line) or Cal520 (labeled at 0.2 uM for 30min). The cells were seeded on fibronectin-coated cover glasses 3 days before experiments. On the day of experiments, the cells were pre-incubated with DMEM with 3mM glucose for 30 minutes either with or without the presence of BMS536924 (2.5uM) or LY294002 (25uM) (Torcis, Avonmouth, Bristol, USA). The cells were then stimulated with various concentrations of either bovine insulin, recombinant human insulin, recombinant human IGF1 (Sigma Chemicals, St. Louis, MO, USA), or glucose.

Fluorescence images of R-GECO or Cal520 were collected and background subtracted. For each trace, fluorescence intensity is normalized to the minimum value before stimulation. P-values are calculated using the Wilcoxon-Mann-Whitney test or student t-test. The type of test to be used in each situation is determined by the P-value computed using the Shapiro-Wilk test to see whether the sample follows a normal distribution. All computations are done using Python 3.7 (code available at https://github.com/yuezheli/primary_cilia_sensitize_insulin_receptor-mediated_negative_feedback).

### Measurement and analysis of membrane potential

MIN6 cells that express primary cilium marker, Arl13b-mScarlet, were seeded on cover glasses 3 days before experiments. Cells were pre-incubated with DMEM with 3mM glucose for 30 minutes before stained with Di-4-ANEQ(F)PTEA (3uM, Potentiometric Probes, Farmington, CT, USA) for 15 minutes. Fluorescence images were collected under either 488 or 640 excitation. Then images went through background subtraction, shade correction, and then emission ratio images under 488 and 640 excitations were generated and quantified based on the plasma membrane labeled by the Arl13b-mScarlet. P values were calculated using Student t test.

## Results

### Insulin alone fails to promote insulin secretion in MIN6 cells

To investigate how insulin feedback specifically affects pancreatic beta cells, we lentivirally transduced mouse insulinoma MIN6 cells with R-GECO (Zhao et al., 2011), a red fluorescence emitting genetically-encoded calcium indicator, to gauge the change of cytosolic calcium in response to glucose. The transduced MIN6 cells were glucose and serum-starved prior to stimulation. When the cells were stimulated with glucose, we observed an increase in R-GECO intensity (Fig 1A, B), suggesting an increase in cytosolic calcium. Statistical analysis shows that the calcium elevation was statistically significant (Fig 1C). As a control for perfusion artifact, stimulating MIN6 cells with low glucose solution did not show a statistically significant change in cytosolic calcium (Fig 1F-H). Additionally, we demonstrated insulin secretion by expressing a GFP-tagged insulin in MIN6 cells. When insulin was secreted, insulin-containing granules generated a sudden increase of fluorescence intensity before vanishing due to diffusion (Fig 1D). The increase of fluorescence intensity was due to a pH change from the acidic secretory granule to the neutral pH environment outside the cell. On average, we observed 5.8 (±1.48, n=15 cells) insulin granules being secreted per minute (Fig 1D). Thus, MIN6 cells elevate calcium and trigger insulin secretion in response to glucose.

**Figure 1.**
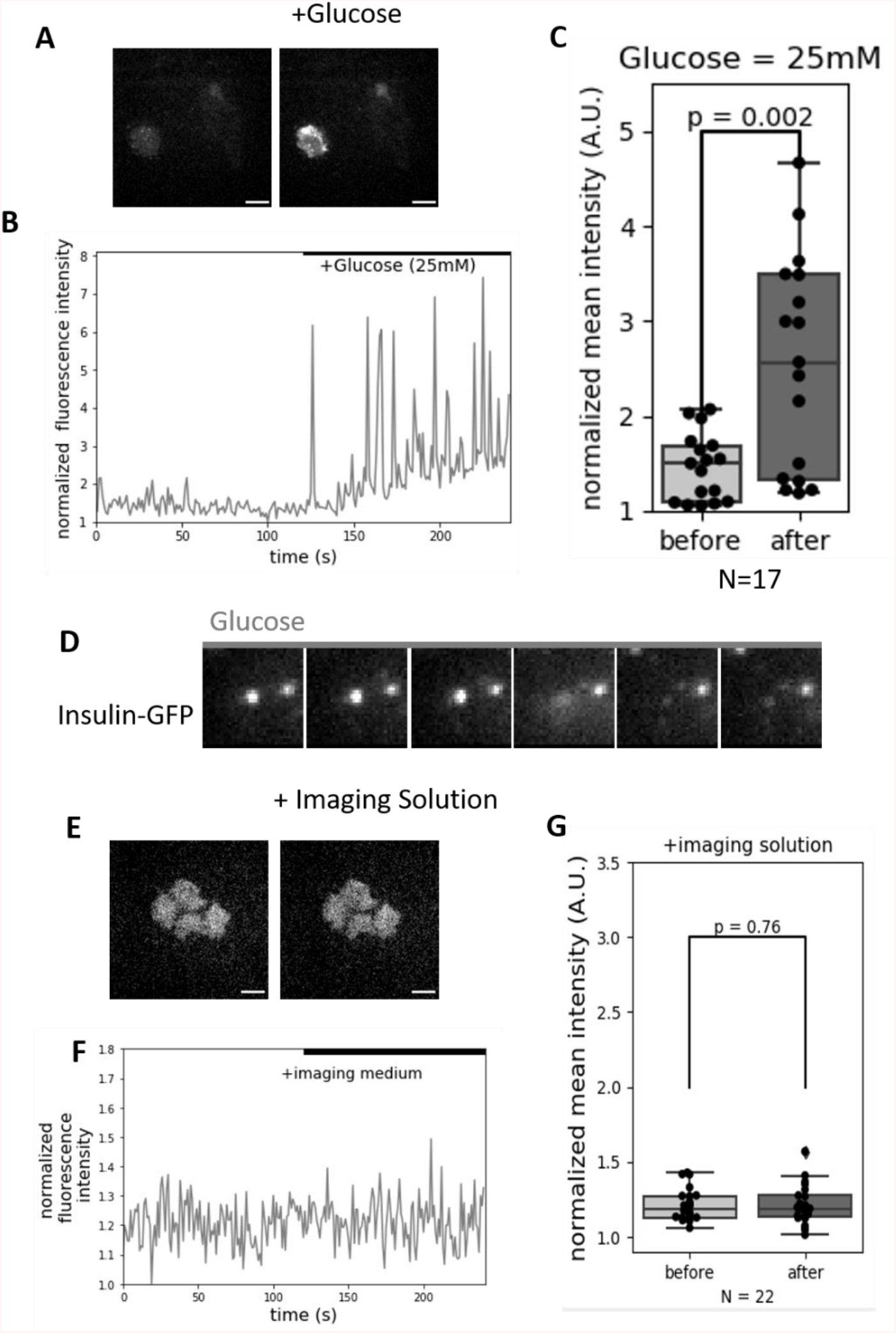
Glucose induced calcium elevation and insulin secretion in MIN6 cells. MIN6 cells that stably express the R-GECO sensor were treated with glucose. The example sensor images before and after treatment (A) and a plot of the normalized fluorescence intensities in time (B) are shown. The quantifications of normalized mean fluorescence intensities are shown in (C). As a control, addition of imaging solution was used to address any potential noise induced by perfusion (E-G). MIN6 cells that express insulin-GFP were treated with 16.7mM glucose (D). A cropped set of sequential images at 1.04s time intervals are shown. See main text for quantifications.

We further validated that insulin receptor (IR) signaling was attenuated after glucose and serum starvation, and insulin triggered IR signaling. Using the phosphorylation level of insulin receptors to gauge the strength of IR signaling, we found a ~50% reduction of IR phosphorylation (Supp. Fig 1A, left 2 lanes) after 30 minutes of glucose and serum starvation; however, prolonged (up to 24 hours) glucose and serum starvation did not further attenuate IR signaling (Supp. Fig 1A, right 3 lanes; Supp. Fig 1B),. To determine if treatment of with either bovine or recombinant human insulin elevates IR phosphorylation, MIN6 cells were treated with 175nM of either source of insulin, both of which displayed ~100% induction of IR phosphorylation (Supp. Fig 1C, first 3 lanes; Supp. Fig 1D). In contrast, blocking insulin signaling with an IR inhibitor, BMS536924, abolished IR phosphorylation (Supp. Fig 1C, last lane; Supp. Fig 1D), suggesting the presence of autocrine feedback of IR activation in MIN6 cells.

Since insulin is known to trigger cytosolic calcium elevation and exocytosis (Aoyagi et al., 2010, Aspinwall et al., 1999, Aspinwall et al., 2000, Hisanaga et al., 2009), we investigated whether exogenous insulin can impact insulin secretion by measuring how cytosolic calcium changes post-insulin stimulation. While cells exhibited different basal states among experiments, we found no statistically significant changes in cytosolic calcium concentration after stimulation with human recombinant insulin at concentrations from 0.1nM to 3.5uM (Supp Fig 2A-C). To investigate the inconsistencies between previous reports and our results, we validated that the insulin source used in our study can induce IR phosphorylation at a concentration as low as 0.1nM (Supp. Fig. 1E). In the concentration range of 0.1nM to 70nM, we found ~20%-40% IR phosphorylation induction (Supp. Fig. 1F). Increasing the insulin concentration also increased phospho-IR concentration: 65% induction after 175nM insulin stimulation, 137% induction after 3.5uM insulin stimulation (Supp. Fig. 1F). 175nM IGF1 treatment led to dramatic induction (~184%). Conversely, we repeated the published experiments (Aoyagi et al., 2010, Aspinwall et al., 1999, Aspinwall et al., 2000, Hisanaga et al., 2009), using >100 nM (175nM) commercially purified bovine insulin to stimulate MIN6 cells. We detected bovine insulin-induced calcium elevation following treatment (Fig 2A, B, upper two rows; Fig 2C, left and middle panel).This contrasted our observations in cells treated with recombinant human insulin (Supp Fig 2A, B, 6th row; Supp Fig 2C, lower second left panel). Western blot analyses revealed 175nM concentrations of recombinant human insulin and bovine insulin induces similar levels of IR phosphorylation (Supp. Fig 1C, D). We were also able to validate that inhibiting PI3K attenuates bovine insulin-induced calcium elevation (Fig 2A-B, bottom panel; Fig 2C, right panel), as reported in Aspinwall et al., 2000. However, the induction was not blocked by the IR inhibitor BMS536924, suggesting bovine insulin triggers calcium elevation in an insulin-independent way.

**Figure 2.**
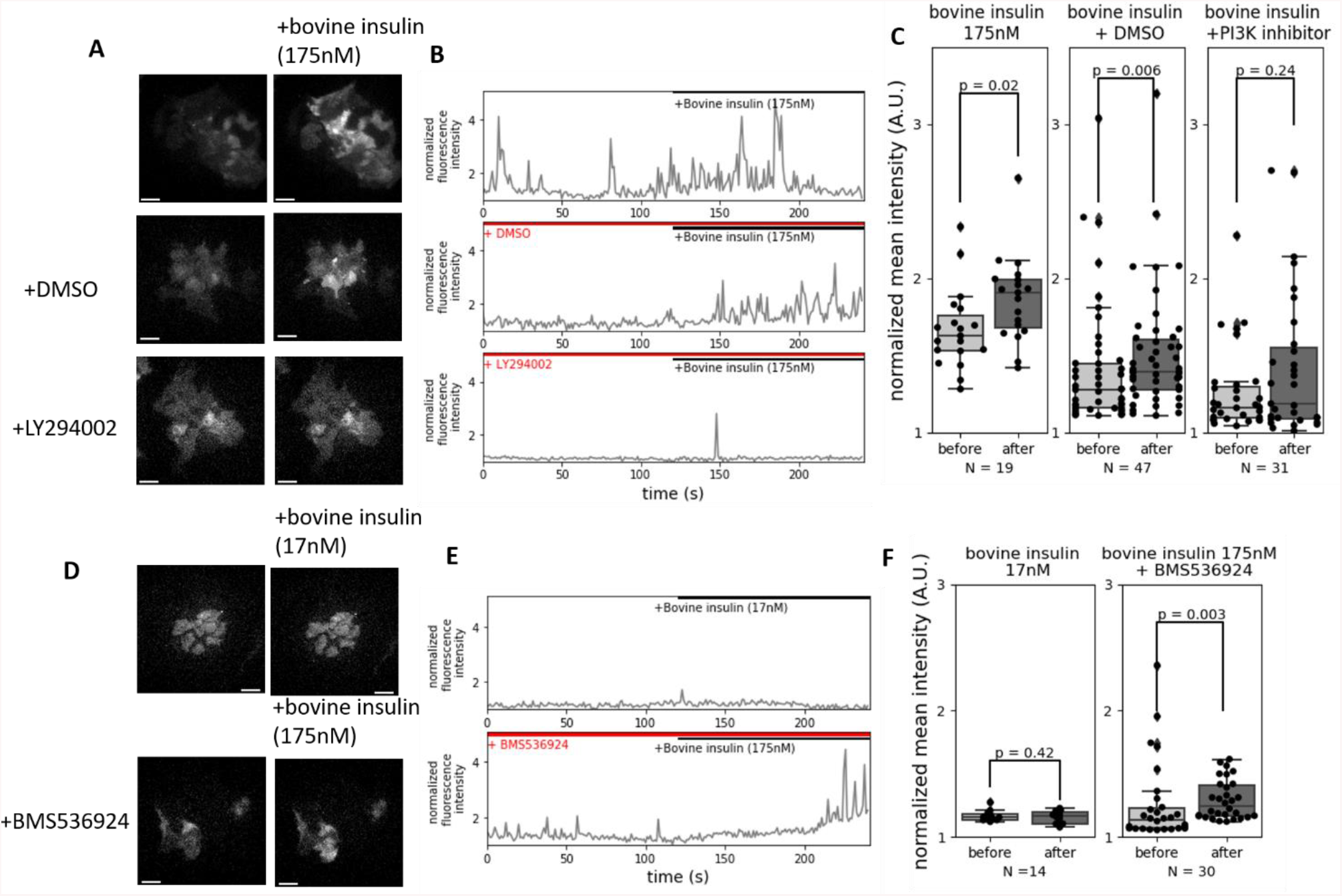
Bovine insulin from a commercial source induced PI3K-dependent, but IR-independent calcium elevation in MIN6 cells. MIN6 cells that stably express the R-GECO sensor were treated with purified bovine insulin in the presence or absence of PI3K inhibitor, LY294002 or DMSO solvent control. The example sensor images (A), plots of normalized fluorescence intensities (B) and their quantifications (C) are shown. Similar experiments were conducted with two different concentrations of insulin and the IR inhibitor BMS536924 (D-F).

### Insulin receptor signaling inhibits glucose-stimulated insulin secretion

Because of the limited brightness of R-GECO sensor, we could only track the calcium changes for no more than 2 minutes after insulin stimulation. Longer or more frequent sensor acquisition often led to increased photobleaching or phototoxicity. Thus, we switched to Cal520, a calcium-sensitive dye, to better track how MIN6 cells may respond to glucose. Consistent with the data obtained from the R-GECO sensor, we observed rapid calcium elevation after glucose addition, reaching its peak values around 2 minutes after glucose addition (Fig 3A, top panel). The calcium elevation gradually attenuated but remained elevated above the initial values for the remainder of the extended 10 minutes of acquisition period (Fig 3B, darker gray bars). Interestingly, co-stimulation of MIN6 cells with glucose and a high concentration (3.5uM) of insulin drastically reduced glucose-stimulated calcium elevation (Fig 3A, second top panel; Fig 3B), in contrast to the positive feedback of insulin reported previously (Aoyagi et al., 2010, Aspinwall et al., 2000, Hisanaga et al., 2009). The insulin-induced inhibition appeared to be insensitive to the diluted concentrations of insulin produced by the MIN6 cells in culture, because inhibiting the basal level of IR signaling failed to have any apparent impact on the amplitude or response time of glucose-stimulated calcium elevation (Fig 3A, lower 2 panels; Fig 3C). Further control experiments showed that BMS536924 alone did not alter the initial cytosolic calcium concentration, regardless cells are in low glucose environment or in high glucose environment (Fig. 3D). Thus, these data argue that, for the observed negative feedback to be physiologically relevant, mechanisms that can sensitize the insulin receptor or enrich the insulin ligand would be required *in vivo*.

**Figure 3.**
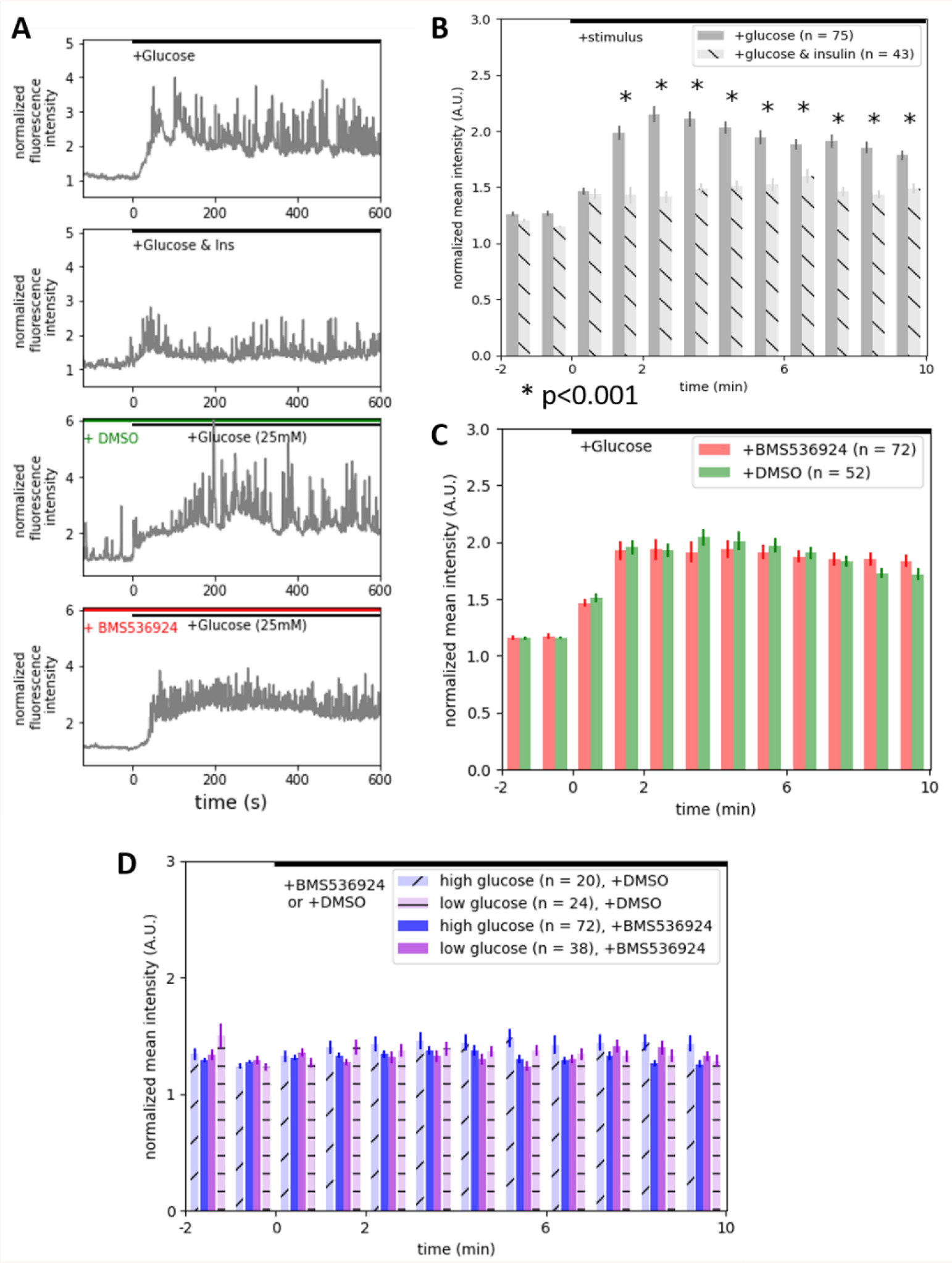
Human recombinant insulin suppressed glucose-induced calcium elevation in MIN6 cells. Cal520 calcium dye was used to label MIN6 cells. The cells were treated with glucose in the presence of absence of human recombinant insulin or IR inhibitor BMS536924. The example trajectories (A) and quantifications of the normalized mean intensities are shown (B and C). Similar experiments were conducted by treating glucose-starved or not starved MIN6 cells with IR inhibitor, BMS536924, or DMSO solvent control (D). Error bars are standard errors.

### Primary cilia promote IR activation and suppress glucose-stimulated calcium elevation

Primary cilia are known to sensitize cell surface receptors. It is reported that most, if not all the pancreatic β cells are ciliated *in vivo* (Gan et al., 2017). However, in MIN6 cells, only ~20-40% of cells are ciliated (Supp. Fig 3G, top panel; Supp. Fig 3H, left bar). Thus, it is possible that ciliated MIN6 cells might have responded to the endogenous insulin, but we failed to detect the effect in bulk measurement due to their small number. To delineate whether primary cilium has a role in glucose-stimulated calcium elevation, we distinguished ciliated cells from cilium-free cells through the expression of Arl13b-mScarlet, a primary cilium marker, in MIN6 cells. By separating the calcium elevation data of ciliated cells from those of cilium-free cells, we found a near 50% reduction of glucose-stimulated calcium elevation in ciliated cells (Fig. 4A-C, Video 1). The cilium-free cells remained sensitive to exogenous insulin and showed suppressed response to co-stimulation of glucose, but no further reduction could be detected in ciliated cells upon insulin stimulation (Fig. 4D-F). Conversely, only ciliated cells responded in calcium elevation to the IR inhibitor, BMS536924, and recovered to the level of cilia-free cells (Fig 4G-I). These data suggested that primary cilia sensitize MIN6 cells to low insulin concentrations and negatively impact cells’ response to glucose.

**Figure 4.**
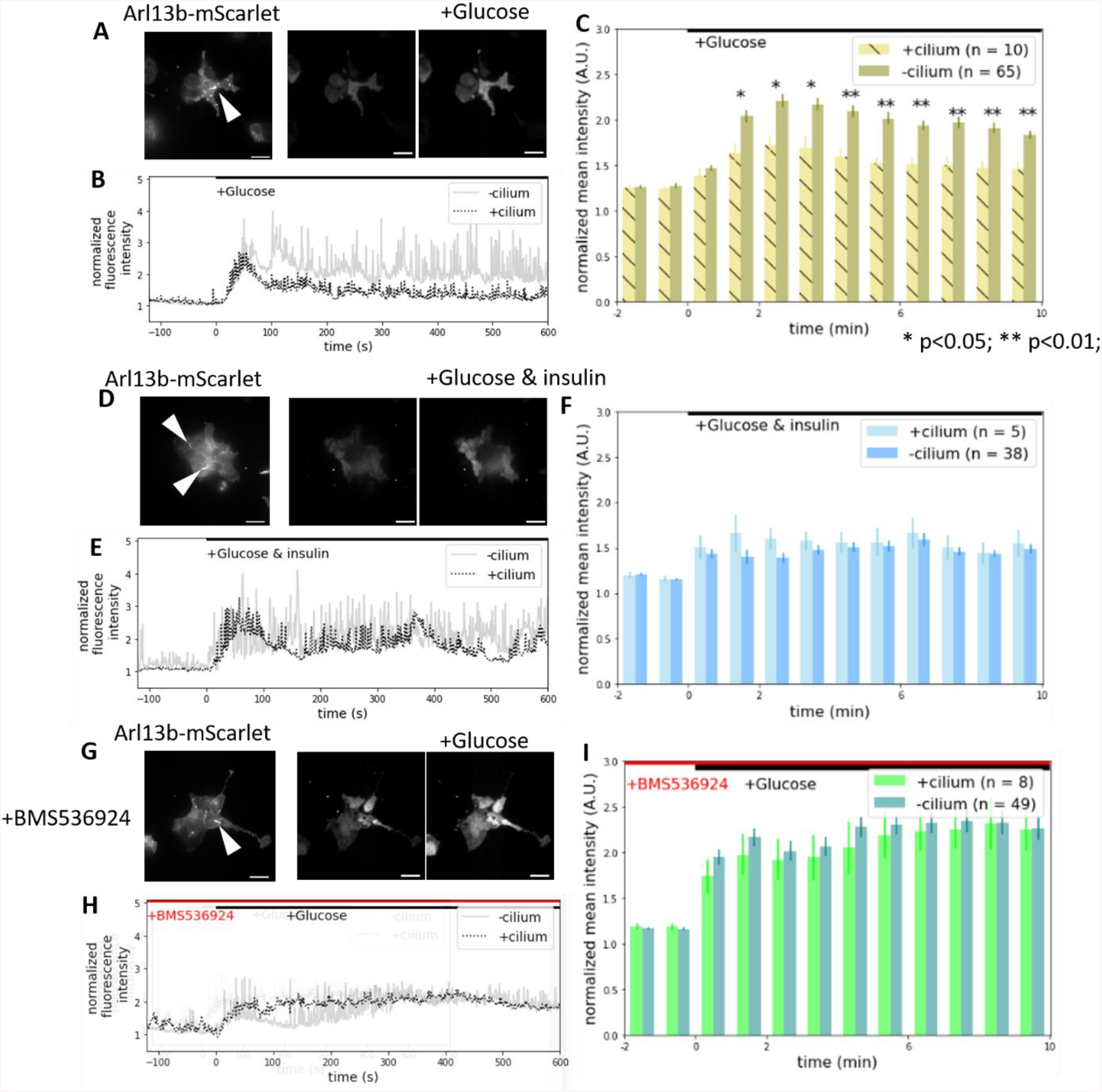
Primary cilia suppressed glucose-induced calcium elevation in an IR-dependent manner. MIN6 cells stably expressing Arl13b-mScarlet, a primary cilium marker, were labeled with Cal520. The cells were treated with glucose alone (A-C), or glucose and insulin (D-F). The example fluorescence images (A, D), Cal520 intensity trajectories (B, E), and quantifications that compared between ciliated and cilium-free cells (C, F) are shown. Similar experiments were conducted by pretreating the cells with the IR inhibitor BMS536924 before adding glucose (G-I). Primary cilia are pointed out with white arrowheads. Scale bar = 20um. Error bars are standard errors.

**Video 1.**
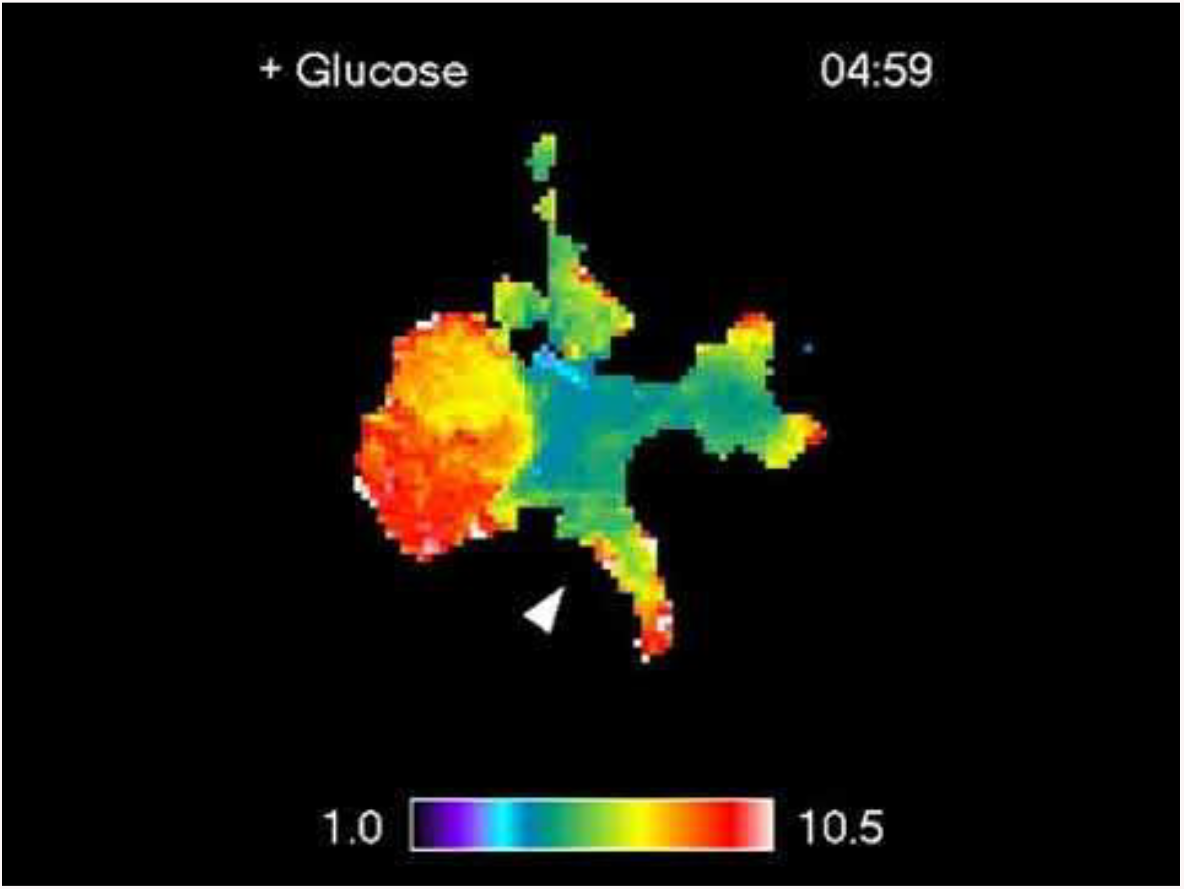
Ciliated MIN6 cell showed attenuated glucose-induced calcium elevation. MIN6 cells stably expressing Arl13b, a primary cilium marker, were labeled with Cal520. The cells were treated with glucose alone. Colormap displays the changes in cytosolic calcium concentration. Ciliated cell is pointed out with white arrowheads.

To examine whether IR is activated in the primary cilia of MIN6 cells, we probed for phosphorylation of the endogenous IR receptor. Cilia-free cells responded to insulin with a significant increase of IR phosphorylation as expected (Fig 5C, left 2 bars). In contrast, we observed strong phosphorylation of IR in primary cilia even in serum-starved cells (Fig 5A, top panel). Adding exogenous insulin had little impact on the ciliary phospho-IR signal, suggesting that the receptor is already highly activated in the primary cilia (Fig 5B, left 2 bars). To further support this idea, we treated cells with sodium orthovanadate, a phosphatase inhibitor (Fig 5D) to saturate the IR phosphorylation. A significant increase of IR phosphorylation in the cilium-free area (Fig 5E) could be observed. However, ciliary IR phosphorylation showed only a minor increase and was not statistically significant (Fig 5F). In addition to the high levels of IR phosphorylation in the primary cilia, the ciliary IR receptor was resistant to the IR inhibitor, BMS536924 (Fig 5A, bottom panel). We found that phospho-IR signal was reduced but remained prominent in the primary cilia, while the phospho-IR signal was completely eliminated outside of cilia or in cilia-free cells (Fig 5B, first and last bar; Fig 5C, first and last bar). These data demonstrated that ciliary IR is highly phosphorylated and is resistant to receptor dephosphorylation.

**Figure 5.**
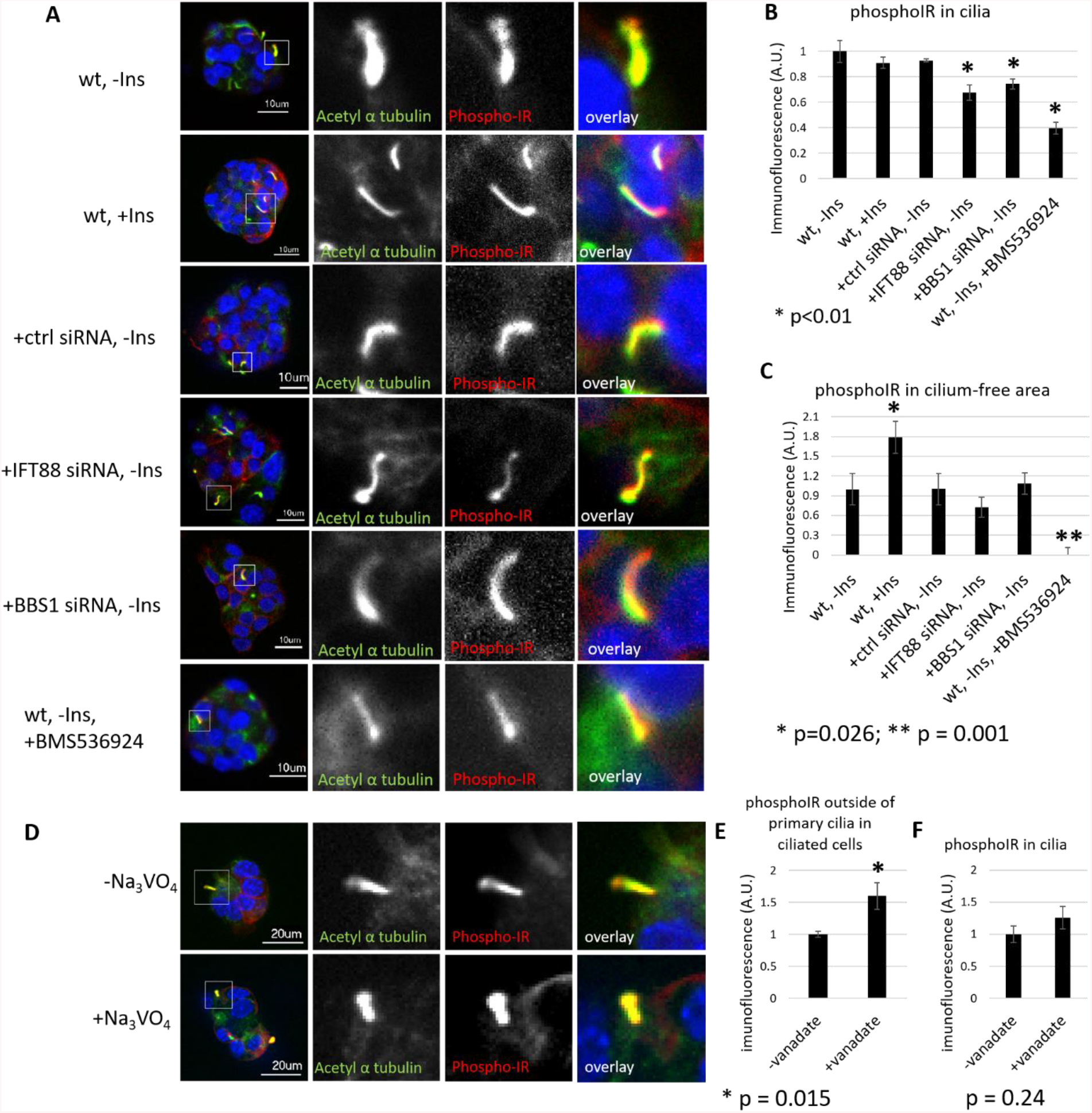
Elevated phosphorylation of IR in the primary cilia and its dependence on ciliary trafficking. (A) MIN6 cells that were treated with or without insulin (top), were transfected with siRNA against IFT88, BBS1 or non-targeting control (middle), or were treated with the IR inhibitor BMS536924 (bottom) were immunostained for acetyl alpha-tubulin, a cilium marker, and phospho-IR. The quantifications of phospho-IR intensities in the primary cilia (B) and in cilium-free area (C) are shown. (D) Similar experiments were performed on MIN6 cells treated with or without a tyrosine phosphatase inhibitor, Na3VO4. The quantifications of phospho-IR intensities in the cilia (F) or outside of cilia of the ciliated cells (E) are shown. Error bars are standard errors.

### Ciliary trafficking is required for IR activation and its suppression of glucose-stimulated calcium elevation

To examine if disrupting ciliary trafficking changes IR phosphorylation, we used siRNA to knockdown IFT88 or BBS1, two proteins known to be involved in ciliary protein trafficking (Starks et al., 2015, Malicki and Avidor-Reiss, 2014, Taschner et al., 2011). A knockdown of these two genes of more than 50% at the protein level was confirmed by western blot analysis (Supp. Fig 3A-D). Consistent with the critical role of IFT88 in cilium assembly (Kim et al., 2010), we observed a reduction of fraction of ciliated cells in IFT88-knockdown cells as demonstrated by co-staining 2 primary cilium markers, acetylated α-tubulin, and Arl13b (Supp. Fig. 3G, H). BBS1-knockdown did not significantly alter the fraction of ciliated cells, but it is known to disrupt ciliary function (Malicki and Avidor-Reiss, 2014). Knockdown of either IFT88 or BBS1, resulted in a decrease in density of phosphorylated IR in primary cilium (Fig 5A, 3rd-5th row, Fig 5B), with no impact on IR phosphorylation outside of primary cilium (Fig 5C). Therefore, defects in IR trafficking may lead to less IR being trafficked into primary cilium, contributing to a decrease in phospho-IR density in the primary cilia.

To determine whether knocking down IFT88 or BBS1 affects IR trafficking in or out of the primary cilia, we ectopically expressed a GFP-tagged constitutively active insulin receptor A (ca IR-A-GFP). Consistent with a previous study (Gerdes et al., 2014), we found that ca IR-A-GFP was localized in the primary cilia in siRNA control cells (Supp. Fig 3I, upper panel; Fig 3J, left bar). However, knocking down either IFT88 or BBS1 blocked ciliary ca IR-A-GFP localization (Supp. Fig 3I, middle and lower panels; Sup. Fig 3J, middle and right bars), demonstrating that both IFT88 and BBS1 are important for ciliary IR trafficking.

To investigate the functional significance of knocking down IFT88 and BBS1, we examined how these knockdown cells elevate calcium in response to glucose stimulation. To reduce cell heterogeneity and siRNA-associated artifact, we continued to rely on the ciliary marker to directly compare between ciliated and cilium-free cells in the same culture. In siRNA control cells, we continued to observe a reduced response to glucose stimulation in ciliated cells compared to cilium-free cells (Fig 6A, B, top panels, Fig 6C). However, knocking down either BBS1 or IFT88 completely abolished the difference in calcium elevation between ciliated and cilium-free cells (Fig 6A, B, middle two panels; Fig 6D, E). In addition to perturbations of ciliary trafficking, we also used siRNA to specifically knockdown IR. Knocking down IR also eliminated the difference in calcium elevation between ciliated and cilium-free cells (Fig 6A, B, bottom panel; Fig. 6F; knockdown validation in Supp. Fig. 3E, F). Thus, these results support that ciliary IR signaling suppresses cells’ response to glucose stimulation.

**Figure 6.**
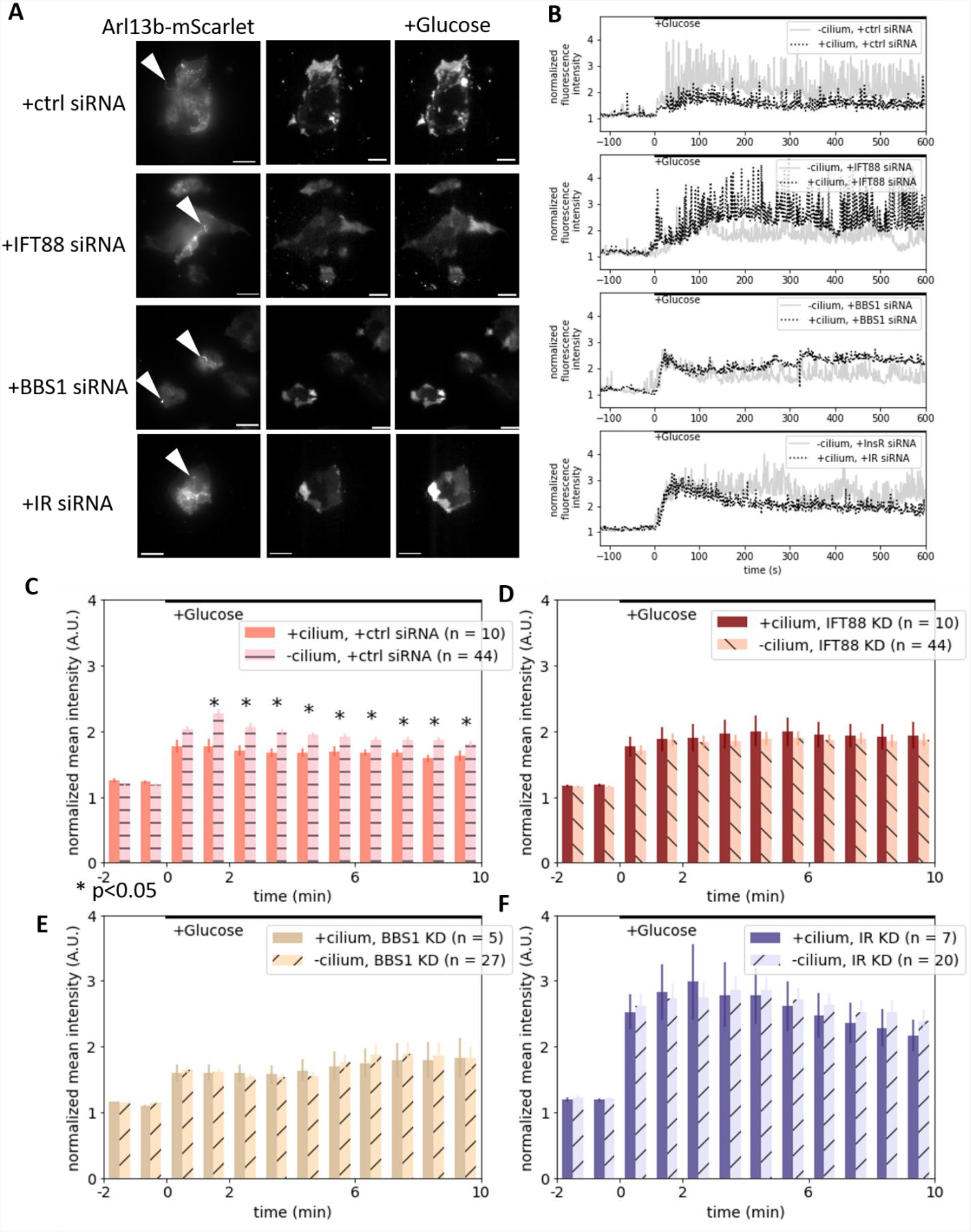
Knockdown of either genes related to ciliary trafficking or IR abolished the inhibition on glucose-induced calcium elevation in ciliated cells. MIN6 cells stably expressing Arl13b-mScarlet were transfected with siRNA control, or siRNA against IFT88, BBS1 or IR. The cells were labeled with Cal520 and treated with glucose as described. The example images (A), and Cal520 intensity trajectories (B), and their quantifications divided between ciliated and cilium-free cells (C-F) are shown. Primary cilia are pointed out with white arrowheads. Scale bar = 20um. Error bars are standard errors.

### Ciliary IR signaling hyperpolarizes the plasma membrane of ciliated beta cells

To investigate the molecular mechanism on how primary cilium asserts a negative impact on GSIS, we evaluated two potential drives behind this cilium-dependent suppression on glucose-stimulated calcium elevation: glucose metabolism, and membrane potential. We used PercevalHR, a biosensor that fluorescence intensity positively correlates with ATP:ADP ratio (Tantama et al., 2013). Upon glucose stimulation, we failed to observe any difference in the fluorescence intensity change of PercevalHR in ciliated and cilium-free cells (Fig 7A; Fig 7B), suggesting ciliary IR signaling has limited impact on glucose metabolism. Since IR signaling was shown to induce membrane hyperpolarization, and thereby inhibit insulin secretion in β cells (Khan et al., 2001), we used a voltage-sensitive dye (Yan et al., 2012) to compare the membrane potentials between ciliated and cilium-free cells. The emission ratio between 488 and 640 excitations is positively correlated with cells’ membrane potential. In control experiments, we stimulated MIN6 cells with tetraethylammonium (TEA), a potassium channel blocker and known to depolarize the plasma membrane (Ni et al., 2010), and observed a ~5% increase in the 488/ 640 ratio (Supp. Fig 4A, B). To avoid interference from electronical coupling between adjacent cells (Speier et al., 2007, Benninger et al., 2008), we only quantified cilium-free cells that were not adjacent to ciliated cells. We observed that the 488/ 640 ratio in ciliated cells were ~10% lower than cilium-free cells (Fig 7C, D). Furthermore, the difference was abolished by treating the cells with the IR inhibitor BMS536924 (Fig 7D right bars). All these results indicate that ciliary IR signaling at least hyperpolarizes plasma membrane of β cells.

**Figure 7.**
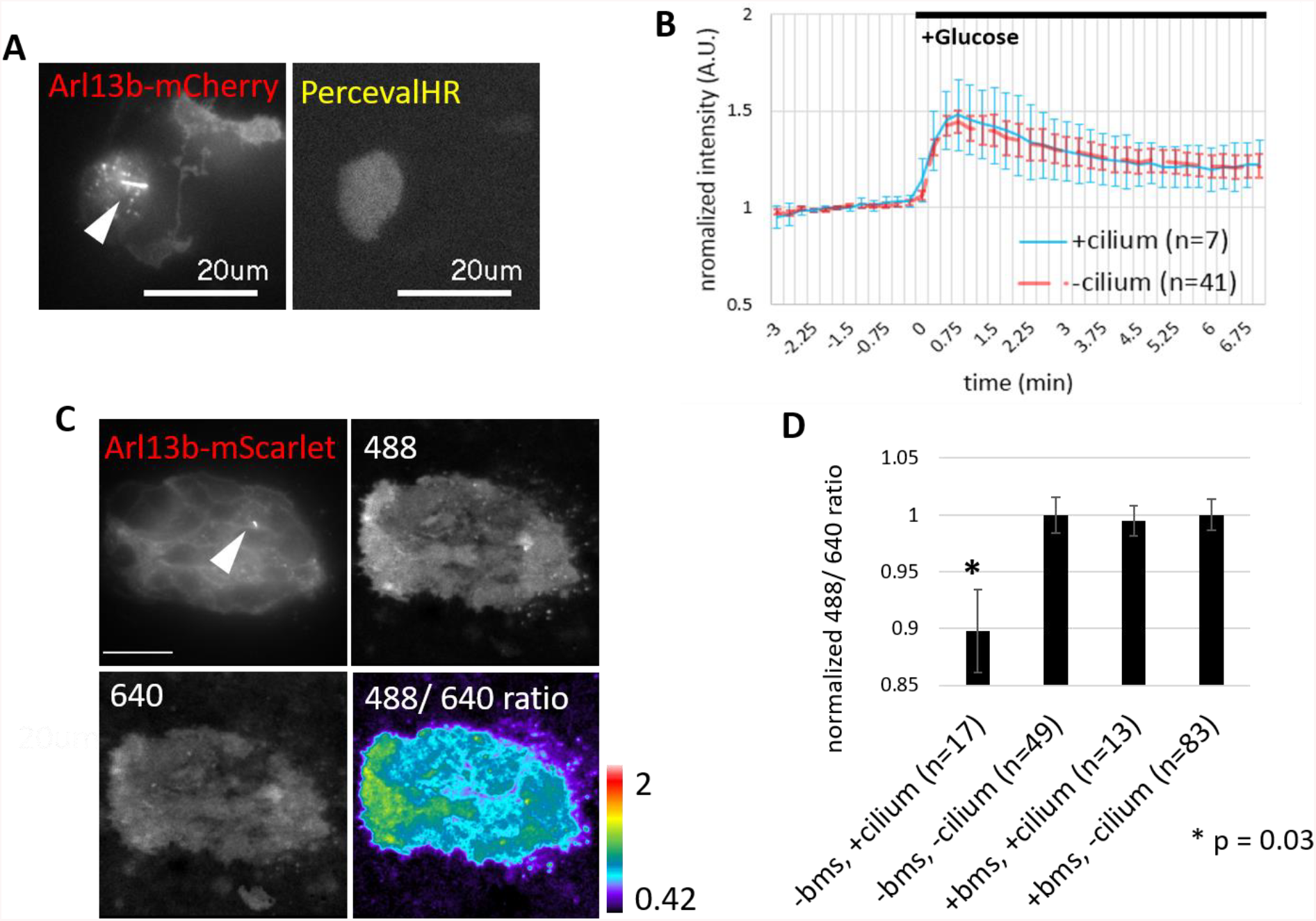
Ciliary IR signaling hyperpolarize the plasma membrane without affecting ATP production. MIN6 cells were co-transfected with Arl13b-mCherry, a primary cilium marker, and PercevaHR, a biosensor for ATP:ADP ratio. The example images (A) and the quantifications of PercevaHR sensor data (B) are shown. (C) MIN6 cells stably expressing Arl13b-mScarlet were labeled with Di-4-ANEQ(F)PTEA, a voltage-sensitive dye. The cells were treated with or without the IR inhibitor BMS536924. The example images (C) and the quantifications of 488/640 ratios (D) are shown. Primary cilia are pointed out with white arrowheads. Error bars in B are 95% confidence intervals, in D are standard errors.

## Discussion

In this study, we revisited the historically opposing observations that were made on insulin feedback in pancreatic β cells. We showed that insulin receptor signaling suppresses glucose-induced calcium elevation in insulinoma MIN6 cells. The negative effect of insulin was reported previously by others (Elahi et al., 1982, Argoud et al., 1987, Persaud et al., 2002, Zhang et al., 1999, Jimenez-Feltstrom et al., 2004). On average, cells stimulated with glucose show a 100% induction calcium elevation, but those that are stimulated with glucose and insulin at the same time have only ~50% of the induction. We also showed that insulin alone has little impact on cytosolic calcium concentration. This is in contrast to what was reported in the literature that bovine insulin increases cytosolic calcium concentration and leads to exocytosis (Roper et al., 2002, Bouche et al., 2010, Aoyagi et al., 2010, Aspinwall et al., 1999, Aspinwall et al., 2000, Hisanaga et al., 2009, Ramsey et al., 2006). We showed these observations are likely due to reagent artifacts from crude bovine insulin. We speculate that the artifacts could be caused by other contaminating molecules secreted by cells in pancreatic islets, such as pituitary adenylate cyclase-activating polypeptide, vasoactive intestinal peptide, and glucagon-like peptide-1 (GLP-1), which are known to potentiate GSIS (Di Cairano et al., 2015, Dickson and Finlayson, 2008, Xin et al., 2016, Segerstolpe et al., 2016, Winzell and Ahren, 2007, Yamada et al., 2004). While the precise concentrations of insulin that β cells expose to *in vivo* are unknown, we examined in this study a broad range of concentrations of insulin on how they affect receptor phosphorylation. We observed insulin-induced insulin receptor phosphorylation plateaued between 0.1nM and 70nM of exogenous insulin stimulation, but unexpectedly, a further increase in exogenous insulin concentration led to a further increase in IR phosphorylation. IGF1 even imposed a more dramatic induction of insulin receptor phosphorylation. We speculate that this increase in insulin/ IGF-induced receptor phosphorylation could be driven by IR -IGF1R hybrid receptors (Benyoucef et al., 2007). These hybrid receptors have a lower affinity to insulin compared to insulin receptors but have a higher affinity of IGF1. The unexpected increase in phospho-IR signal could be due to our phospho-IR antibody that can pick up phospho-IGF1R as well.

Our observation that ciliated cells have less GSIS appears to be in contrast to the report in the literature that both BBS4 −/− and Tg737 ORPK mice have impaired GSIS (Gerdes et al., 2014, Zhang et al., 2004). We saw similar impacts of disrupting primary cilia or their function in siRNA experiments (Fig 5 C-E), but the focus of the study was on specifically the ciliary IR instead of the primary cilia compartment in general. We speculate that ciliopathy mouse models have impaired GSIS is more likely due to the long-term effect of malfunction of primary cilia. For example, the disruption of cell polarity (Walz, 2017, Boehlke et al., 2010) may disrupt the proper release of insulin, as most of the secretion takes places towards the vasculature (Low et al., 2014) or into the interstitial space (Takahashi et al., 2002). Alternatively, primary cilia may also modulate GSIS by altering the energy homeostasis either through other receptors (Volta et al., 2019) or other parts of an organism that can influence pancreatic β cells (e.g. brain, adipose tissue) (Oh et al., 2015).

We found that primary cilia potentiate insulin receptor phosphorylation and suppress calcium elevation in the absence of exogenous insulin. The sensitized ciliary IR phosphorylation could be due to a lack of phosphatase activity inside the primary cilia because ciliary IR phosphorylation appeared to resist the treatment of IR or phosphatase inhibitor. The elevated IR signaling from primary cilium could also be the result of cilia’s high surface-to-volume ratio, an environment that promotes access of the activated IR to concentrated or specific downstream targets. This phenomenon is not new, as Zhu et al., 2009 reported that ciliary IGF1R is sensitized to ligand stimulation. However, it is not clear how ciliary IR signaling propagates from primary cilia to the rest of the cells. One possibility is that ciliary IR signaling increases ATP-sensitive potassium channels’ overall conductance through a PI3K-dependent pathway, resulting in suppressed GSIS. This is consistent with our observation, and literature suggests that IR signaling, in general, increases the overall conductance of ATP-sensitive potassium channels in a PI3K-dependent manner. (Khan et al., 2001, Zaika et al., 2016, Xu et al., 2015). This is also consistent with the observation in Gerdes et al., 2014 that disrupting primary cilia resulted in an altered PI3K and Akt phosphorylation. However, the caveat of this idea is that the downstream effectors, IRS-1, does not go into primary cilia. It is reported that IRS-1, phospho-IRS-1, Akt, and phospho-Akt are detected at the basal bodies, adjacent but outside the primary cilia (Zhu et al., 2009). Thus, it is not clear how IR inside primary cilia phosphorylates IRS-1 outside primary cilia. Alternatively, ciliary IR may signal by cross-talks between receptor tyrosine kinases and G-protein coupled receptors (Gavi et al., 2006). This may enable primary cilia to signal through cAMP to modulate β cell physiology.

*In vivo*, primary cilia are localized at the apical surface of beta cells, exposed to an extracellular lumen that is sealed off by tight junctions and is distant from vasculature (Gan et al., 2017). Thus, the exact insulin concentration that the beta cells are exposed to *in vivo* is difficult to predict. We speculate this extracellular lumen may have a lower insulin concentration and may be less subject to transient fluctuations due to the presence of surrounding tight junctions. It is likely that the negative feedback effect we uncovered in this study is related to the homeostasis of beta cells. The attenuation of the calcium elevation may provide a tonic inhibition to prevent beta cells from burnout and promoting β cell survival (Bae et al., 2019, Yoshimura et al., 2010, Luciani et al., 2013, Prentki and Nolan, 2006).

## Author’s contribution

YW and YL conceive the project. YL and PKS validate the cell line. YL characterizes MIN6 cells’ response to different stimuli under different conditions, ciliary insulin receptor signaling and enrichment, and membrane potential difference between ciliated and cilium-free cells. YW and YL write the manuscript.

## Acknowledgments

We thank Dr. C. Acker for his help in imaging membrane potential using voltage-sensitive dye, Di-4-ANEQ(F)PTEA. We thank Dr. L. Wilson for critical reading and editing of the manuscript. This work has been supported by N.I.H. grants GM117061 (Y.I.W.)

## Supplementary Data

**Supplementary Figure 1.**
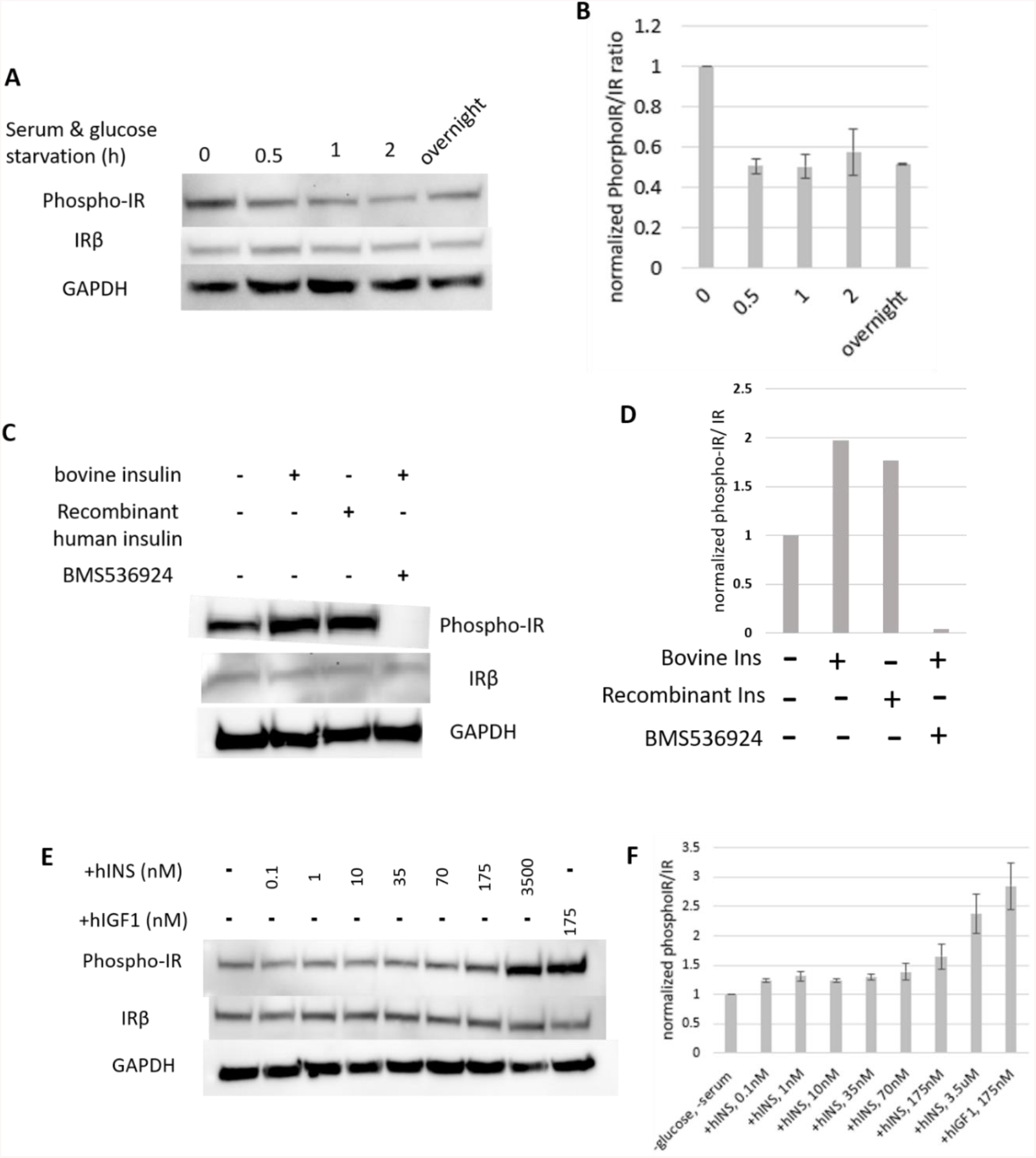
Characterization of insulin receptor phosphorylation in MIN6 cells. At basal level, insulin receptor was phosphorylated in MIN6 cells (A, first lane; B, first bar). With half an hour glucose and serum starvation, basal insulin receptor phosphorylation dropped by half (A, second lane; B, second bar). Prolonged glucose and serum starvation did not further decrease basal insulin receptor phosphorylation (A, third to last lane; B, third to last bar). Treatment of 175nM of either bovine or recombinant human insulin elevated insulin receptor phosphorylation by a fold (C, first to third lane; D, first to third bar). Pretreating MIN6 cells with 2.5uM of BMS536924, an insulin receptor/ insulin-like growth factor 1 receptor inhibitor, efficiently inhibited insulin receptor phosphorylation (C, last lane; D, last bar). Furthermore, different concentrations of insulin stimulation, ranging from 0.1nM to 3.5uM, all led to IR phosphorylation (E), with the maximum strength of over a fold of induction (F). IGF1 led to more significant IR phosphorylation.

**Supplementary Figure 2.**
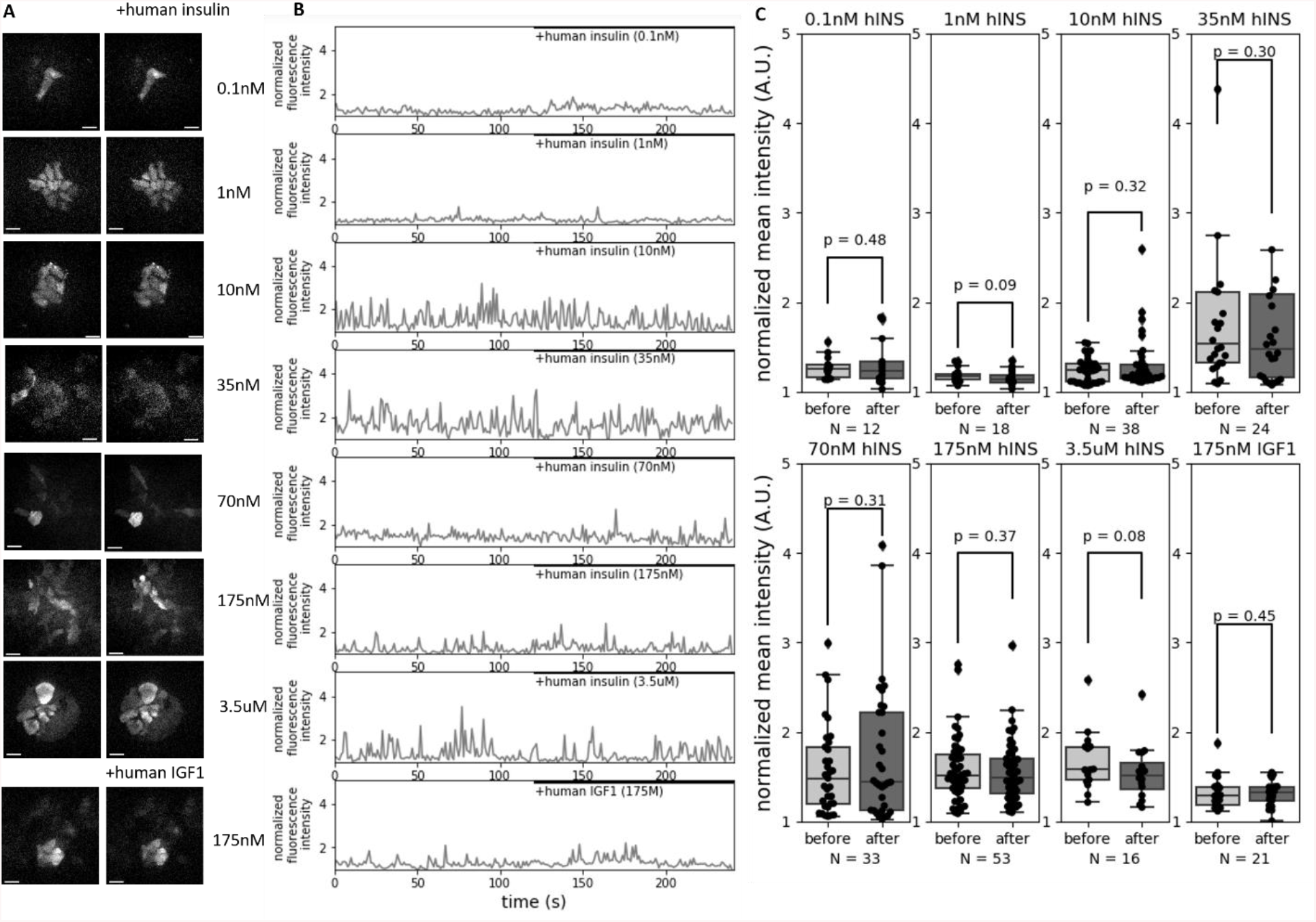
Recombinant human insulin or recombinant human IGF1 did not change cytosolic calcium concentration. Recombinant human insulin, with concentration ranging from 0.1nM to 3.5uM, showed no impact on cytosolic calcium concentration in MIN6 cells (sample images in A top 7 panels; sample traces in B top 7 samples; quantification in C top 4 panels and lower left 3 panels). Recombinant human IGF1 also showed no impact on cytosolic calcium concentration in MIN6 cells (sample images in A bottom panel; sample trace in B bottom panel; quantification in C lower right panel).

**Supplementary Figure 3.**
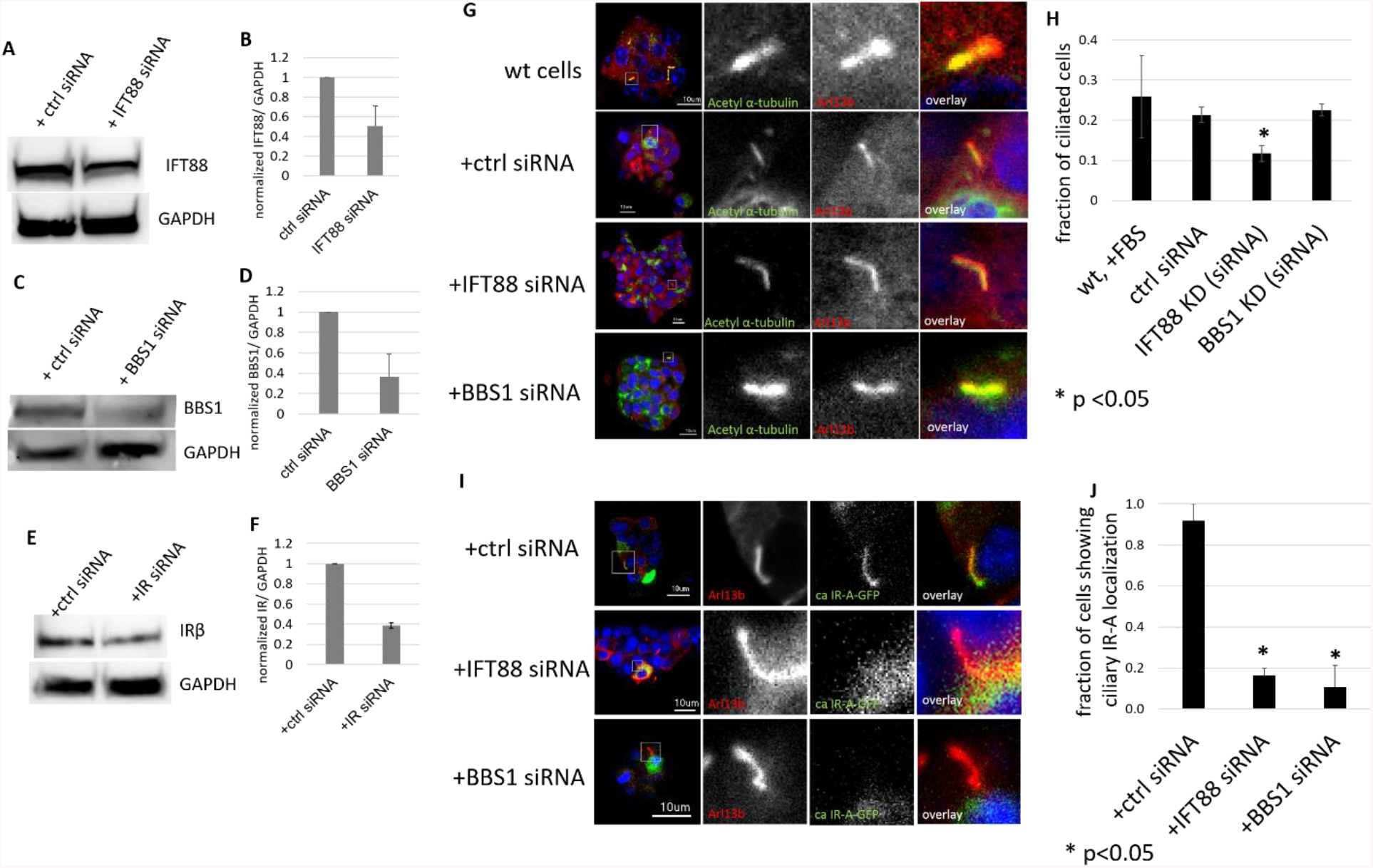
Characterization of siRNA knockdown of IFT88, BBS1 and IR in MIN6 cells. Knocking down of IFT88, BBS1, IR using siRNA were validation using Western blots, respectively (A-F). Overall, there was a ~60% knockdown efficiency for each protein (B, D, F). Cells transfected with non-specific control siRNA showed similar percentage of ciliated cells (sample images in G top 2 panels; quantification in H left 2 bars). IFT88 led to a decreased percentage of MIN6 cells that had a primary cilium (sample image in G 3rd panel; quantification in H 3rd bar), while BBS1 knockdown had no significant impact on the percentage of ciliated cells (sample images in G bottom panel; quantification in H right bar). Constitutive active IR-A (ca IR-A) was enriched in primary cilia (sample images in I top panels; quantification in J left bar), but the knockdown of either IFT88 or BBS1 led to the loss of ciliary IR localization (sample images in in I lower 2 panels; quantification in J right 2 bars).

**Supplementary Figure 4.**
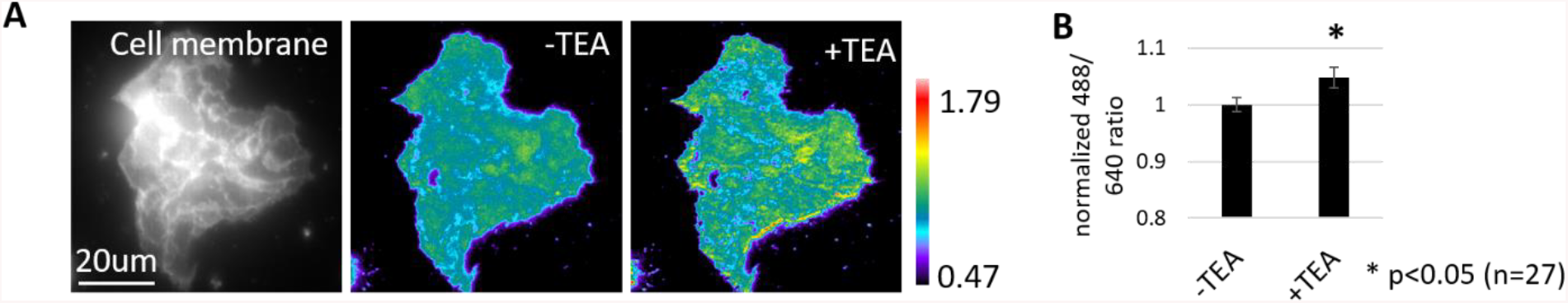
Characterization of membrane potential in MIN6 cells. This is measured by the emission ratio of Di-4-ANEQ(F)PTEA, a voltage-sensitive dye under 488 and 640 excitations. A higher ratio indicates higher membrane potential. Treating MIN6 cells with tetraethylammonium (TEA), a reagent known to depolarize plasma membrane, led to an increase of 488/ 640 ratio (sample image in A, scale bar = 20um; quantification in B).

